# Genetic control of mRNA splicing as a potential mechanism for incomplete penetrance of rare coding variants

**DOI:** 10.1101/2023.01.31.526505

**Authors:** Jonah Einson, Dafni Glinos, Eric Boerwinkle, Peter Castaldi, Dawood Darbar, Mariza de Andrade, Patrick Ellinor, Myriam Fornage, Stacey Gabriel, Soren Germer, Richard Gibbs, Craig P. Hersh, Jill Johnsen, Robert Kaplan, Barbara A. Konkle, Charles Kooperberg, Rami Nassir, Ruth J.F. Loos, Deborah A Meyers, Braxton D. Mitchell, Bruce Psaty, Ramachandran S. Vasan, Stephen S. Rich, Michael Rienstra, Jerome I. Rotter, Aabida Saferali, M. Benjamin Shoemaker, Edwin Silverman, Albert Vernon Smith, NHLBI Trans-Omics for Precision Medicine (TOPMed) Consortium, Pejman Mohammadi, Stephane E. Castel, Ivan Iossifov, Tuuli Lappalainen

## Abstract

Exonic variants present some of the strongest links between genotype and phenotype. However, these variants can have significant inter-individual pathogenicity differences, known as variable penetrance. In this study, we propose a model where genetically controlled mRNA splicing modulates the pathogenicity of exonic variants. By first cataloging exonic inclusion from RNA-seq data in GTEx v8, we find that pathogenic alleles are depleted on highly included exons. Using a large-scale phased WGS data from the TOPMed consortium, we observe that this effect may be driven by common splice-regulatory genetic variants, and that natural selection acts on haplotype configurations that reduce the transcript inclusion of putatively pathogenic variants, especially when limiting to haploinsufficient genes. Finally, we test if this effect may be relevant for autism risk using families from the Simons Simplex Collection, but find that splicing of pathogenic alleles has a penetrance reducing effect here as well. Overall, our results indicate that common splice-regulatory variants may play a role in reducing the damaging effects of rare exonic variants.

## Introduction

Incomplete penetrance is a well known phenomenon, where an individual carries a disease-associated allele, but develops no symptoms of the disease themself (Forrest et al. 2022; Gettler et al. 2021; Shawky 2014). Similarly, variable expressivity refers to analogous gradual differences in disease severity; here we refer to both as variable penetrance. These instances are likely underreported in the literature due to ascertainment bias, when many studies are based on sequencing due to a prior genetic condition (Cooper et al. 2013; Dewey et al. 2016). Even amongst Mendelian disease variants, which are typically thought of as having strong effects on phenotype, differing levels of severity have been observed between carriers (Chen et al. 2016). These changes have been attributed to epistatic or additive effects of genetic modifiers, as well as environmental modifiers of penetrance, which can be difficult to control in an experimental setting (Maya et al. 2018). When looking at incomplete penetrance in specific diseases, genetic modifiers have been mapped, for example, to BRCA in breast cancer (Milne and Antoniou 2011), and RET in Hirschsprung’s disease (Emison et al. 2005). Modified penetrance has also been studied in the context of polygenic risk scores, where multiple common risk variants increase the expected pathogenicity of a disease-relevant variant (Fahed et al. 2020). However, genome-wide patterns underlying modified penetrance are still poorly known. One potential mechanism for incomplete penetrance are cis-regulatory mechanisms that affect the regulation of a gene carrying a pathogenic variant. This model has been tested with expression quantitative trait loci (eQTLs) acting as modifiers of penetrance (Castel et al. 2018), but can be expanded to other types of gene regulatory processes, such as mRNA splicing. While eQTLs control the dosage of their target genes, splicing alters inclusion of variant-carrying exons in transcripts, which could potentially have a large effect on the overall pathogenicity of a damaging variant.

Alternative splicing is responsible for the great diversity of isoform structures observed across human tissues and cell types (Keren et al. 2010). With regard to coding variant interpretation, exons with lower expression have been shown to be less likely to harbor pathogenic variants, while ubiquitously included exons can be prioritized for gene disrupting rare variants (Cummings et al. 2020). Autistic individuals with variants on the same exons have been shown to have remarkably similar disease phenotypes, putatively due to the variants having similar effects on gene dosage or function, a notable finding given the extreme heterogeneity of the condition (Chiang et al. 2021). Additionally, splicing can be influenced by common genetic variation, as evidenced by the many studies that use large scale WGS and transcriptomic datasets to map splicing quantitative trait loci (sQTLs) (Alasoo et al. 2019; Consortium 2020; Garrido-Martín et al. 2021; Kerimov et al. 2020). sQTLs in general have been implicated in disease risk and other genetic traits (Li et al. 2016; Noble et al. 2020; Ongen and Dermitzakis 2015).

In this study, we build upon the finding that transcript usage of genes containing alleles contributes to the allele’s pathogenicity, and ask if common splice-regulatory variants may partially drive this phenomenon and affect inter-individual variation in penetrance. Expanding on previous methodology (Castel et al. 2018), we look for non-random haplotype combinations of sQTL variants and putatively pathogenic rare variants in population scale datasets. Such an observation could indicate that haplotype combinations have an effect on fitness, and by proxy, disease risk. In doing so, we develop a general framework for modeling common and rare variant haplotypes in a population, with a corresponding test to detect deviations from the null (Figure 1, Supplemental Figure 1). These analyses will improve our understanding of how variants across the annotation and allele frequency spectrum act together to shape human traits and could ultimately aid our interpretation of rare variants in a clinical context.

**Figure 1.**
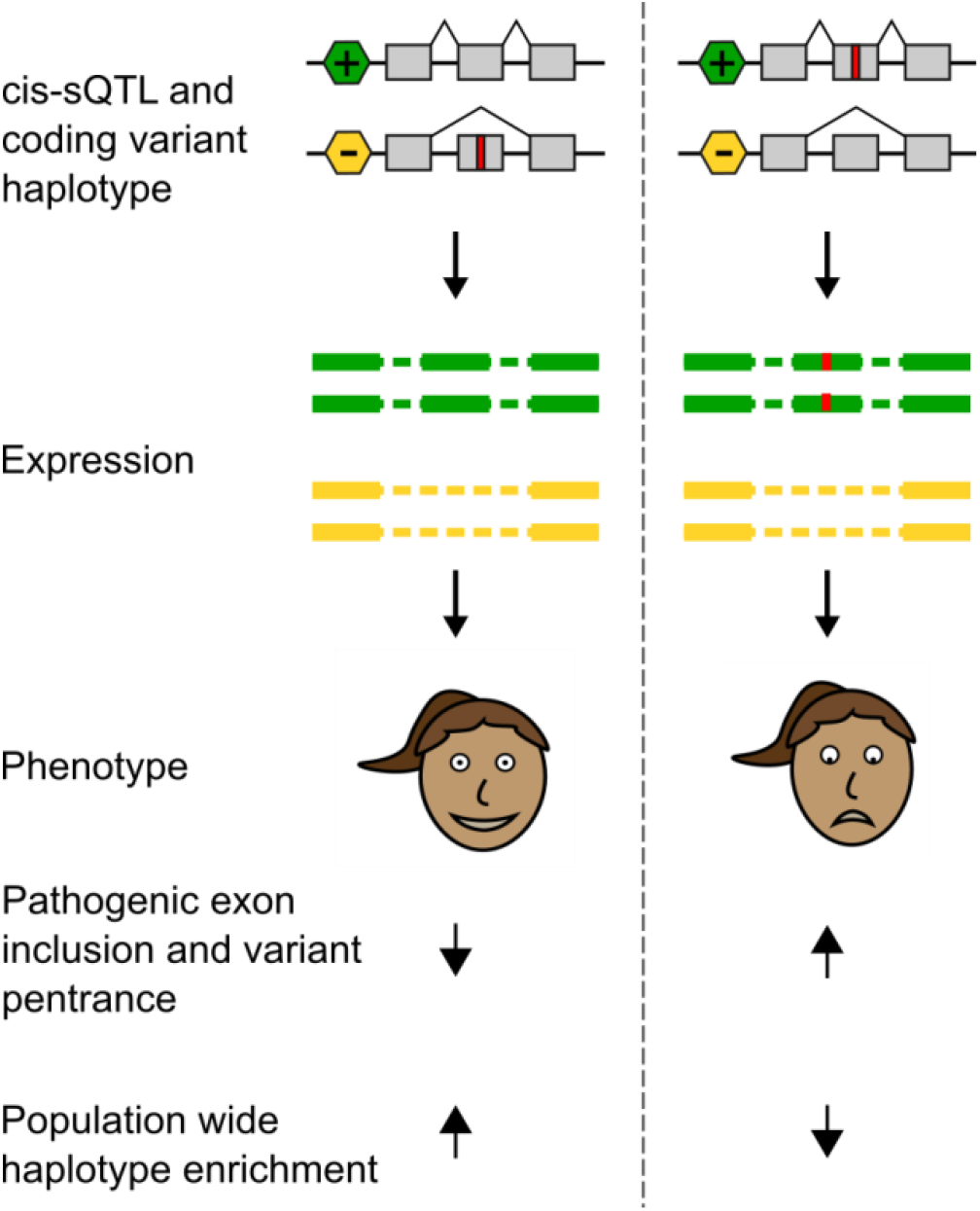
Splice-regulatory variants as modifiers of penetrance hypothesis. The hypothesis of this study is illustrated with an example of an individual who is heterozygous for both a ψQTL and a coding variant. The two possible haplotype configurations result in either a reduced or increased penetrance state of the coding allele, depending if the allele is on the more lowly or highly included exon respectively. We predict that natural selection would deplete those that fall in a high penetrance configuration in the general population. See Supplementary Figure S1 for a quantitative description of the model.

## Results

### Deleterious rare alleles accumulate at lowly spliced exons with respect to the population

We first tested the hypothesis that rare pathogenic alleles (CADD > 15) (Rentzsch et al. 2019) are more likely to occur at less spliced-in exons (Figure 1). To accomplish this, we used bulk RNA-sequencing (RNA-seq) and whole genome sequencing (WGS) data from the Genotype Tissue Expression Project (GTEx) v8 release, which is representative of a general population free of severe genetic disease. We defined variants as rare if their variant frequency in gnomAD (Karczewski et al. 2020) was less than 0.5% and they appeared 5 or fewer times among the 838 GTEx WGS donors.

To begin, we calculated percent spliced in (PSI) scores for all annotated protein-coding gene exons across 18 GTEx tissues, and only kept exons with sufficient splicing variability across individuals (Methods, Supplemental Table 1, Supplemental Figure S2A). We extracted rare alleles that fell on variably spliced exons, separating alleles within 10bp of a splice junction to avoid cases where the allele is more likely to directly affect splicing. To compare the splicing of each donor with a deleterious allele to the population distribution per exon, we calculated PSI Z-scores across all tissues with available data (Supplemental Figure S2B, Methods). We found that PSI Z-scores were significantly different between exons carrying deleterious (N = 19,178) and non-deleterious (N = 49,575) rare alleles (Mann-Whitney U-Test: *p* = 2.577×10^−4^). This rank difference was accounted for by a modest decrease in mean PSI Z-score among donors that carried deleterious alleles in a given exon, which was consistent across tissues and across variant consequence annotations (Figure 2, Supplemental Figure S3). Notably, stop-gained variants had the strongest association with low PSI Z-scores - even stronger than the signal for variants close to splice junction - but the overall result was present for multiple annotation categories (Supplemental Figure S3). This suggests that the signal is not solely driven by the most pathogenic variants nor direct rare variant effects on splicing. These results extend the previous work, comparing different exons and showing accumulation of stop-gained variants on those with lower inclusion (Cummings et al. 2020). Here, observe a similar pattern when comparing different individuals within a given exon, consistent with the hypothesis that the penetrance of coding alleles is reduced when they fall on more lowly included exons. However, this approach does not discern the underlying reasons for splicing differences between individuals, including alleles that may drive a decrease in splicing and their haplotype combinations with rare alleles.

**Figure 2:**
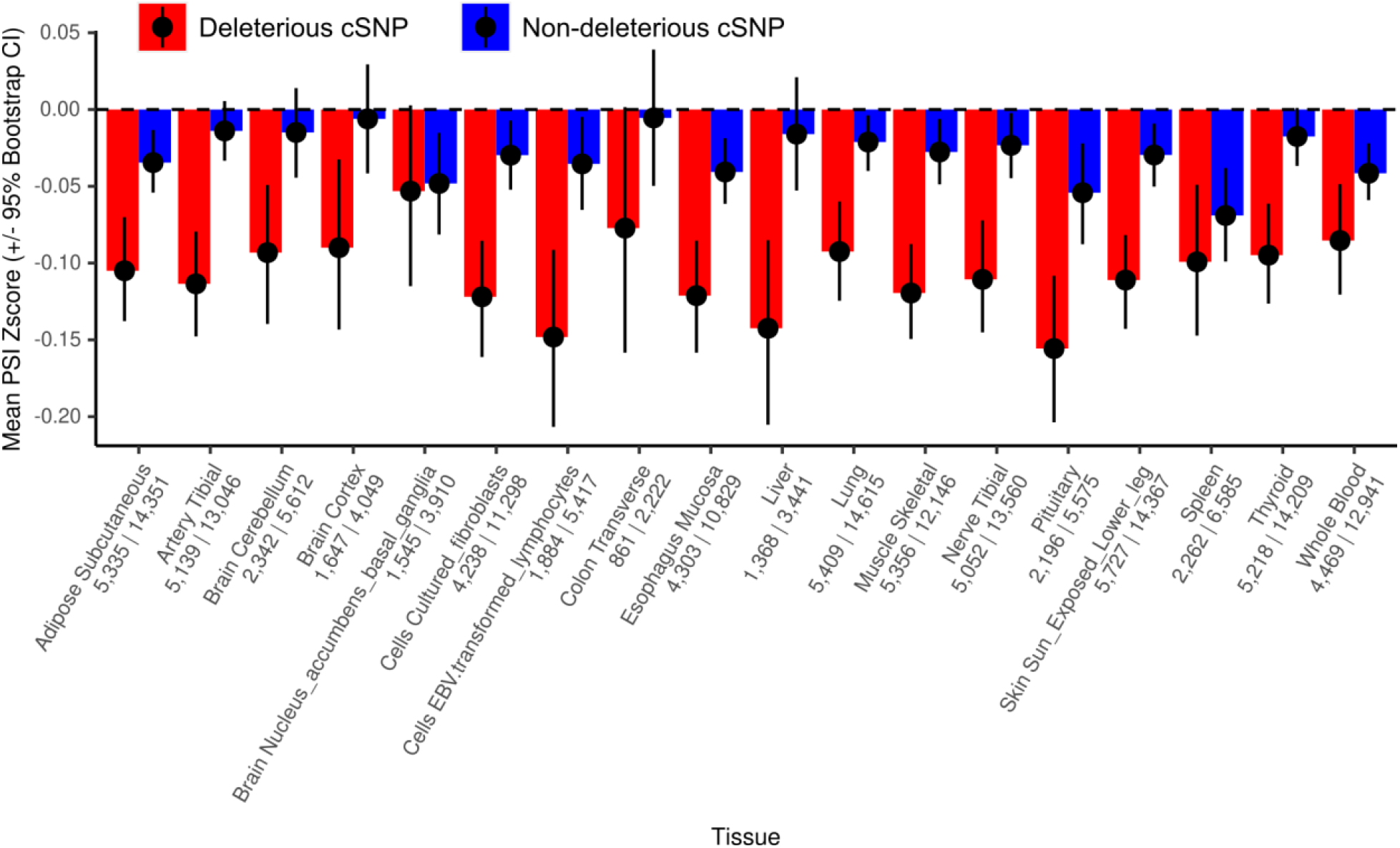
Mean PSI Z-scores across tissues. Mean decrease in PSI Z-scores among individuals carrying rare alleles at variably spliced exons across 18 GTEx tissues, split by deleterious (CADD > 15) and non-deleterious (CADD < 15) rare variants. The number of deleterious and non-deleterious alleles respectively are printed below each tissue name. Error bars represent 95% bootstrapped confidence intervals.

### A general model for coding allele-QTL haplotype configurations

We next sought to test if regulatory alleles on the same haplotype as rare coding alleles contribute to this phenomenon, using phased whole genome sequencing (WGS) data. Since directly quantifying the penetrance of coding alleles is difficult, our approach was to observe modified penetrace through the lens of purifying selection, where high-penetrance haplotype combinations would be depleted from the general population. Advantageously, this technique allows us to use large phased WGS datasets where individual gene expression data is not available.

Initially, splice-regulatory alleles were cataloged in GTEx through quantitative trait locus (QTL) mapping, using the percent spliced in (PSI or ψ) (Pervouchine et al. 2013) of each exon as a quantitative phenotype. These alleles are hence referred to as ψQTLs. We use the “ψ” nomenclature to differentiate from sQTLs, where the splicing phenotype can vary between studies and is often less interpretable for downstream applications. ψQTL mapping and properties are described in (Einson et al. 2022). Briefly, we mapped ψQTLs from GTEx v8 using the same filtered set of PSI scores across 18 tissues as in the previous analyses (see Methods). We compiled a set of 5,196 cross-tissue ψQTL genes (one sVariant and one sExon per gene), and recorded which alleles led to higher or lower sExon inclusion. We also mapped secondary sExons across ψQTL genes where the top sVariant was also associated with splicing in the same direction as the top sExon in the same gene, which were used to expand the amount of genic space where rare variants could be considered.

Next, to robustly test for non-random haplotype combinations of rare exonic alleles and common ψQTL alleles, we describe an approach that quantifies the significance of deviations in haplotype combinations from the null in a dataset, taking variable ψQTL allele frequencies into account:In most datasets, ψQTL alleles that may have an effect on rare variant penetrance are non-uniformly distributed, and thus we expect an unequal number of high and low penetrance haplotypes under the null (Figure 3). To account for this, we model these data using the Poisson-Binomial distribution, a generalization of the Binomial distribution describing the sum of *n* independent but non-identically distributed Bernoulli random variables. (González et al. 2016; Hong 2013; Wang 1993) When looking at counts of haplotype combinations, the probability of observing a high-penetrance haplotype is assigned according to the relevant ψQTL allele frequency, independently across QTL genes. To apply the model to haplotypes extracted from phased genetic data, we developed a bootstrapping procedure that approximates the cumulative distribution function of the Poisson-Binomial, constituting a convenient method for calculating the significance, enrichment/depletion effect sizes (ε) and confidence intervals when comparing enrichment scores between groups i.e. haplotypes with deleterious vs. non-deleterious rare alleles (see Methods for details). In simulations, our method was well powered to detect deviations from the null across all tested theoretical allele frequency distributions, and performed well against other methods that directly calculate and approximate the CDF of the Poisson-binomial. (Figure 4, Supplemental Figure S4). This approach is generalizable to other analyses of haplotype combinations; here we apply it to test nonrandom combinations of ψQTL and rare coding alleles.

**Figure 3:**
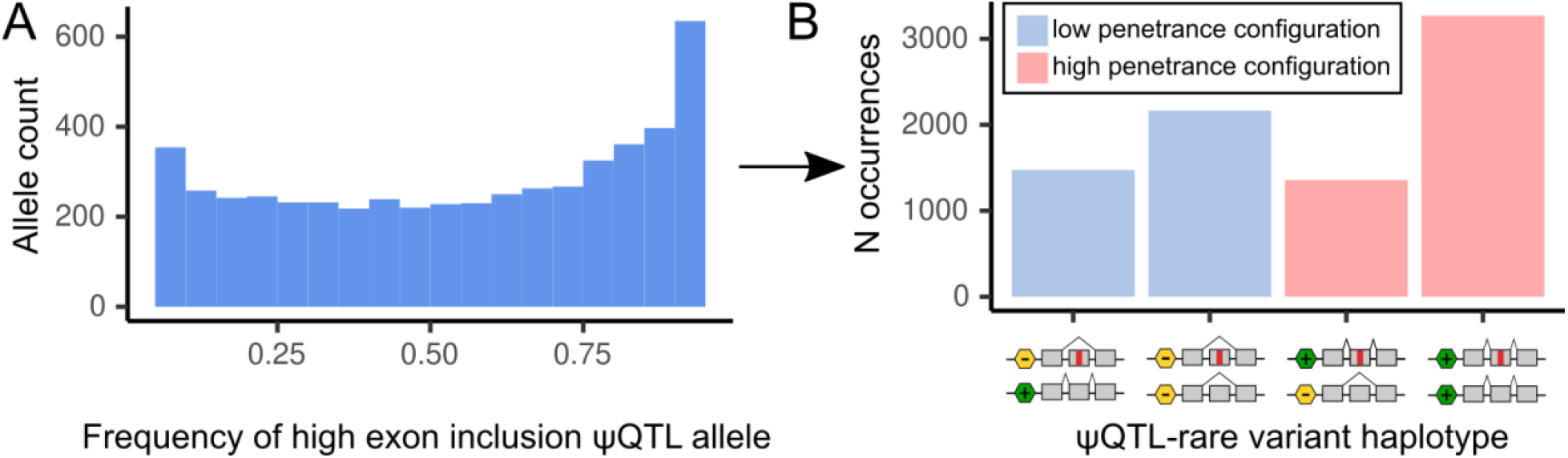
ψQTL high inclusion allele frequencies and haplotype counts in GTEx. A. Distribution of allele frequencies for ψQTLs that lead to higher exon inclusion. High inclusion ψQTL allele frequencies are skewed to the right, meaning ψQTLs that include their target exon are more common in the general population. B. As a result of the nonuniform frequency distribution of high inclusion sQTL alleles, we expect to see more high penetrance haplotype configurations in general. This motivates the necessity to design a test that accounts for this difference.

**Figure 4:**
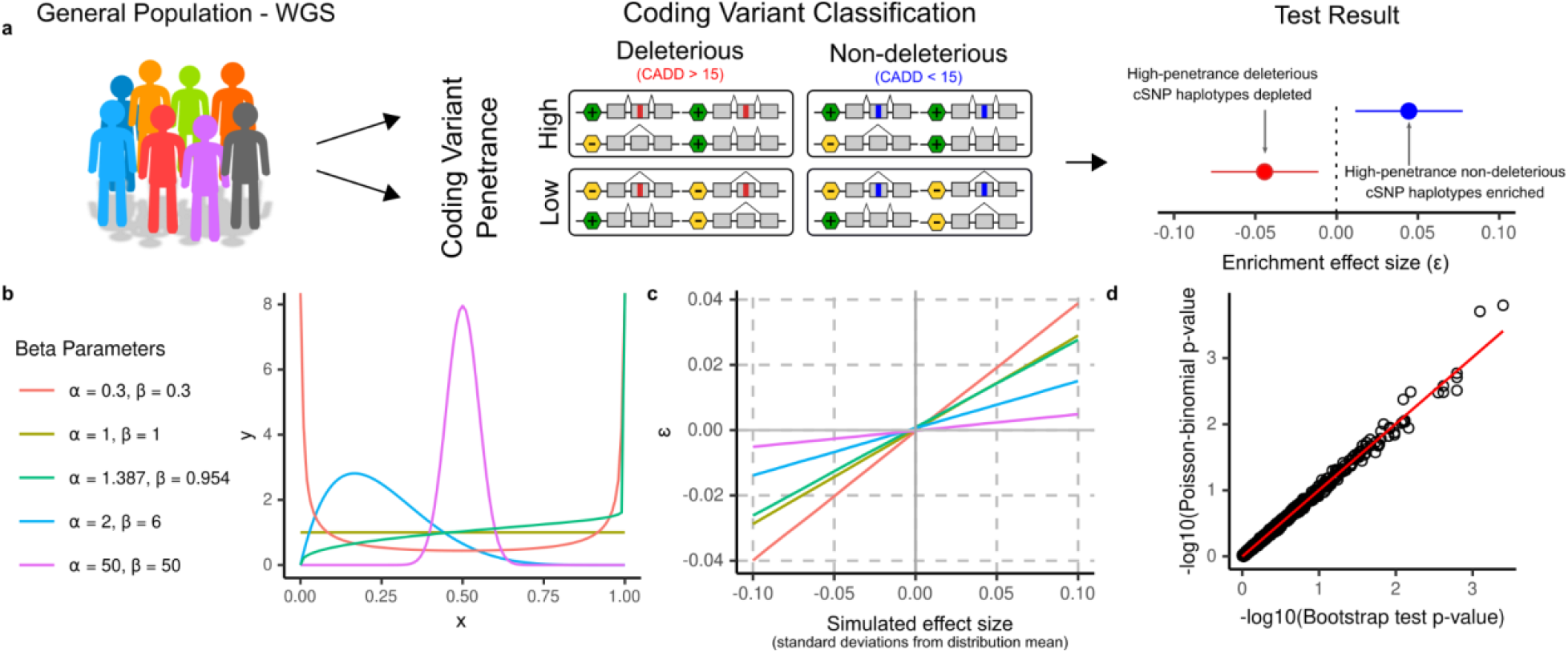
The Poisson-binomial distribution models haplotype configuration counts. **a**. We use phased variant calls from WGS across large populations to test for deviation in the frequencies of ψQTL-coding variant haplotype configurations. The magnitude and effect direction of deviation, which we call ε, is calculated using a procedure described in Methods. The magnitude of ε - but importantly not its direction - depends on the underlying ψQTL allele frequency distribution, as the probability of observing a high penetrance haplotype is dependent on the ψQTL allele frequency at each gene. Counts of highly penetrant haplotypes are modeled by the Poisson-Binomial distribution. When running our test, we frequently divide haplotypes into those with deleterious (CADD > 15) and non-deleterious (CADD < 15) coding variants, which serve as a negative control where we do not expect to see evidence of purifying selection. **b**. To verify that our test captures deviations from the null under any theoretical allele frequency distribution, we simulated datasets by drawing samples from various Beta distributions with different parameters. The Beta is defined by shape parameters α and β. The parameters α = 1.387 and β = 0.954 were estimated from the high-inclusion ψQTL allele frequency distribution in GTEx using the Method of Moments estimator. **c**. We benchmarked our test by simulating data from distributions with increasingly larger deviations from the expected mean, in order to test how the magnitude of ε differs depending on the input distribution. This diagram can be used as a reference for how to interpret the magnitude of epsilon, given a dataset’s underlying probability distribution **d**. P-values from a simulated dataset of haplotypes from 1,000 individuals across 1,000 genes, with ψQTL allele frequencies matching those in GTEx. We find that our method accurately replicates the results from the Poisson-binomial distribution, calculated using the ‘poibin’ (Hong 2013) R package.

### High penetrance haplotypes are depleted in TOPMed and GTEx

After defining a theoretical model that describes counts of common regulatory alleles and rare coding alleles in a given population, we tested three datasets for evidence of selection against high penetrance coding alleles driven by genetically regulated splicing.

#### Enrichment in GTEx

We identified ψQTL-rare allele haplotypes using population and read-backed phased (Castel et al. 2016) WGS data from GTEx V8, labeling haplotypes in putative high and low penetrance configurations according to whether the rare alternative allele was on the higher or lower inclusion ψQTL haplotype, respectively (Figure 1 & 3). We limited our analysis to European-Americans, since the ψQTL data is dominated by European ancestries, with rare variants annotated to potentially deleterious (CADD > 15) and non-deleterious (CADD < 15) variants as described in Methods. In total, 14,767 haplotypes were identified, spanning 714 individuals and 2,475 genes (Supplemental Figure S5). We observed an overall depletion of putative high-penetrance haplotypes (ε = -0.0156, Poisson-binomial test *p* = 1.006×10^−6^), consistent with our hypothesis. However, we did not detect a stronger depletion for putatively deleterious rare alleles (*p* = 0.508, Figure 5), possibly due to the modest sample size of GTEx limiting our statistical power.

**Figure 5:**
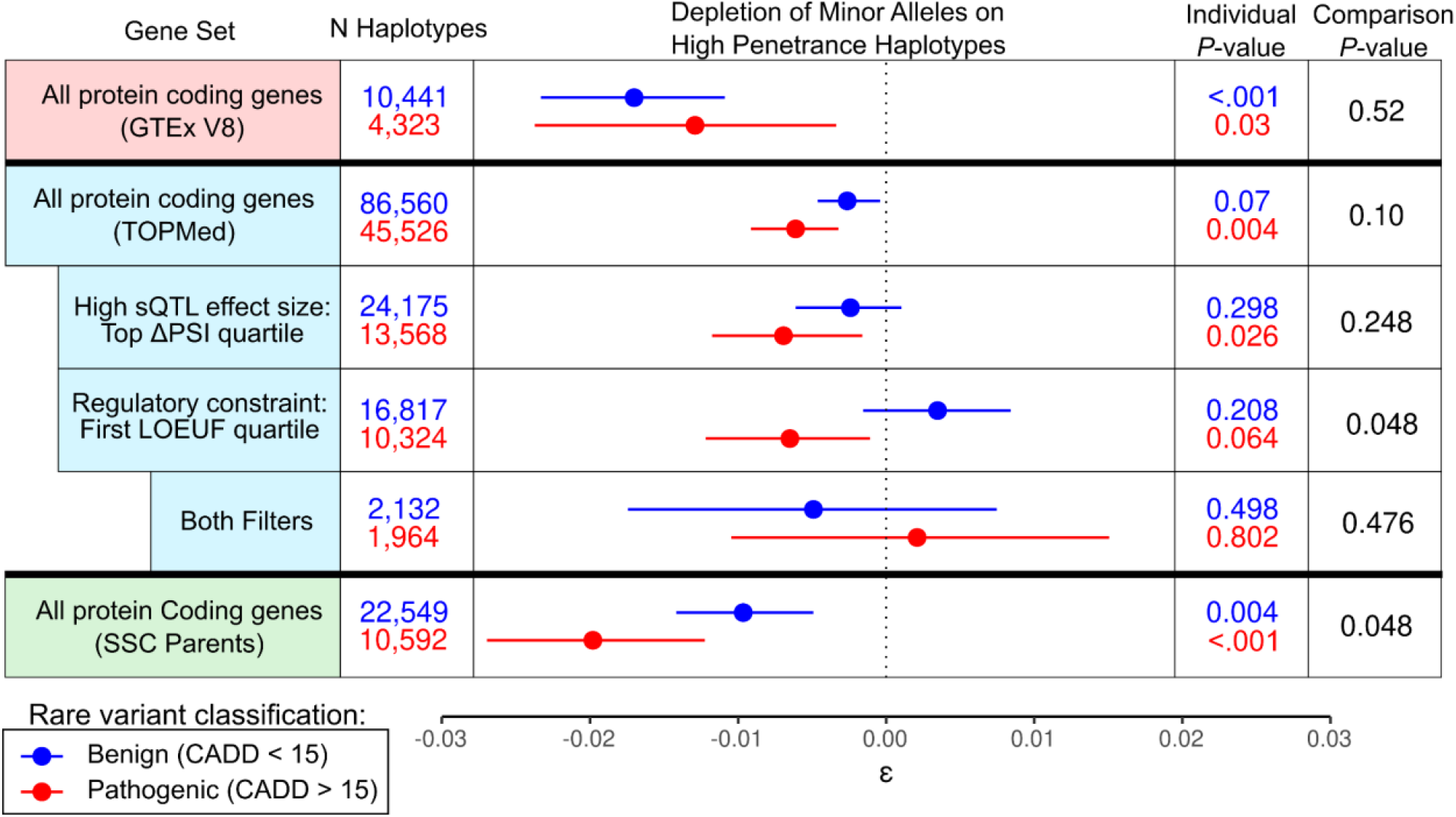
Rare alleles carried in predicted high penetrance ψQTL configurations in GTEx, TOPMed, and SSC Parents. We tested for deviation in the frequencies of coding allele - ψQTL configurations across all protein coding genes with a significant ψQTL. A negative value of ε indicates fewer haplotypes than expected given the population’s ψQTL allele frequencies. Individual *p-*values and 95% confidence intervals were generated using our approximation of the Poisson-binomial cdf, with 1,000 bootstraps. Comparison *P*-values were generated with 1,000 bootstraps.

#### Enrichment in TOPMed

Next, we increased our power to detect evidence of selection against putative high penetrance haplotypes by using population-phased WGS data from 44,634 European-American ancestry individuals in 19 TOPMed cohorts, post-filtering (Methods, Supplemental Figure S5). The large sample size in TOPMed allowed us to limit the analysis to exonic variants with 10 or fewer occurrences (excluding singletons due to limitations of population-based phasing), or <0.0213% minor allele frequency. With the same set of ψQTLs from GTEx, we identified the haplotype of 38,869 rare alleles that fell in primary and secondary sExons. Across all protein-coding genes and rare alleles, we observed a modest but significant overall depletion of high penetrance haplotypes than expected (ε = -0.0037, Poisson-binomial *p* = 3.43×10^−4^). Haplotypes with putatively deleterious rare alleles had some indication of being more depleted than those with non-deleterious rare alleles, but not to a degree that reached statistical significance (*p* = .100, Figure 5). However, we hypothesized that this result would be more pronounced in genes with stronger ψQTLs, as well as genes known to be intolerant to loss of function variation. When focusing on genes with stronger ψQTLs where the ΔPSI score was in the top quartile (ΔPSI > 0.076) the difference was again not significant (*p* = 0.248). However, when quantifying gene constraint with LOEUF (Karczewski et al. 2020) and limiting to genes in the first quartile among sGenes (LOEUF < 0.460), we detected a significant difference in high-penetrance haplotype depletion between the two groups (*p =* 0.048), suggesting that splicing may play a greater role in modifying penetrance in genes known to be constrained. Finally, while we would expect to see the greatest effects of purifying selection among constrained genes with strong ψQTLs, the small number of such genes limits our power and no significant association was detected (*p* = 0.982). We found that across genes in general, ΔPSI and LOEUF were positively correlated, so genes with high ΔPSI and low LOEUF were uncommon (Supplemental Figure S6C). While subtle, these results suggest that deleterious rare alleles are more likely to be carried on exons that are skipped due to the effects of common regulatory variants, especially in constrained genes.

Next, we wanted to explore if any genes or classes of genes drove our observation of high-penetrance haplotype depletion. To this end, using the same TOPMed data, we tested for nonrandom haplotype combinations on a gene-by-gene basis, instead of pooling haplotypes across all genes as in the previous approach. For 2,396 genes with more than 10 ψQTL-coding variant haplotypes across all available individuals, we ran a Poisson-binomial test for high-penetrance haplotype depletion (Supplemental Figure S7). We observed little signal, with approximately equal numbers of genes with enrichment and depletion of high and low penetrance haplotypes. However, only 411 of the genes had more than 30 deleterious allele haplotypes, indicating that our power is quite limited. Thus, our results indicate that observing signals of modified penetrance at the gene level in population cohorts is very challenging.

### Genetically controlled splicing’s contribution to disease gene variant penetrance

In addition to studying the general population as above, we next turned to investigate nonrandom distribution of ψQTL-coding allele haplotypes in a disease cohort: the Simons Simplex Collection (SSC) with 2,380 Autism Spectrum Disorder (ASD) simplex families. Rare coding variants are known to contribute to the etiology of ASD (Iossifov et al. 2014; Sanders et al. 2015; Sanders et al. 2012), and the large set of transmission-resolved WGS data available in the SSC make it a suitable dataset to search for haplotype patterns indicative of modified penetrance. While de novo variants also play an important role in autism risk (Iossifov et al. 2014), their number is so low that we chose to focus on inherited variants.

First, we sought to replicate the depletion of potential high-penetrance haplotypes observed in TOPMed, using SSC parents, who are a cohort of unrelated individuals, phenotypically healthy but with potential enrichment of ASD risk variants due to having a child with ASD. We analyzed all genes with a ψQTL in GTEx, limiting our analysis to coding alleles with 3 or fewer occurrences across all parents, and removing genes with an unusually high number of rare variant haplotypes (Supplemental Figure S5). Singleton variants were included, since their haplotype can be confidently resolved using phasing by transmission. We recapitulated the patterns observed in TOPMed, with a significant depletion of high-penetrance haplotypes with deleterious rare alleles (ε=-0.019, Poisson-binomial *p* = 2.11×10^−8^), with high-penetrance haplotypes carrying deleterious rare alleles more depleted than those carrying non-deleterious rare alleles (Comparison p-value = 0.042, Figure 5).

Next, we sought to analyze potential splicing modifiers of the penetrance of disease-causing alleles in SSC by focusing on rare inherited variants in ASD-implicated genes. These alleles, while potentially contributing to ASD in the proband, are also carried on the same haplotypic background by a healthy parent and often a healthy sibling. Thus, both increased or decreased penetrance ψQTL configurations could be possible (Supplemental Figure S8) To test this, we analyzed deviation in haplotype frequencies in parents, probands, and siblings, among the 218 out of the 1,010 genes implicated in ASD risk according to SFARI Gene (Banerjee-Basu and Packer) that also had a ψQTL. No significant deviation was detected in SSC parents (ε = - 0.0278, *p =* 0.122). Interestingly, across probands and unaffected siblings we found that putatively highly penetrant haplotypes with deleterious coding alleles were depleted (ε = -0.055 & -0.047, *p* = 0.020 & 0.088 respectively). While it seems counterintuitive to see depletion of penetrant haplotypes in individuals with ASD, we reason that this penetrance reducing effect may be acting to protect parents from developing phenotypes of ASD. We find that the SFARI genes tend to be highly constrained, compared to all protein coding genes (Supplemental Figure S8B) (Neale et al. 2012), and that these same alleles were also highly depleted among unrelated individuals in TOPMed (Figure 6), further corroborating the overall observed pattern of selection for penetrance reducing haplotype combinations.

**Figure 6:**
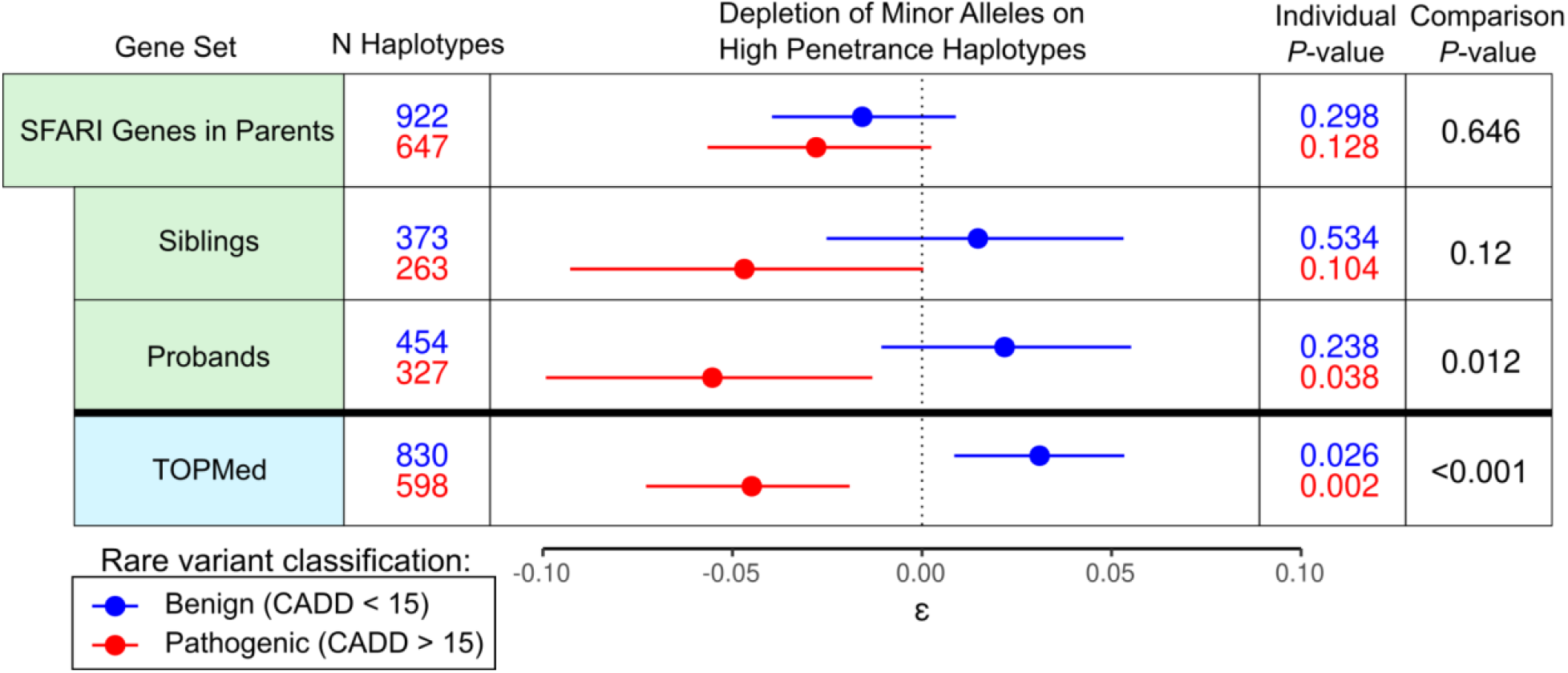
ψQTL haplotype configurations in Autism Spectrum Disorder implicated genes in ASD families. We tested for deviation in the frequencies of high penetrance variant - ψQTL configurations in ASD-implicated genes in parents, probands and unaffected siblings in SSC families.

## Discussion

In this study, we have expanded our model of *cis*-regulatory alleles as modifiers of penetrance of coding variants (Castel et al. 2018) to directly consider splice-regulatory alleles as potential additional drivers. We first show that individuals carrying potentially deleterious rare mutations at variably spliced exons tend to use those exons in transcripts less frequently. This observation could indicate that the penetrance of these rare alleles is reduced by their exclusion from transcripts. However, this approach does not reveal the reason. One approach to potentially shed light on this would be analysis of allele-specific transcript structure, but this is not possible with short read RNA-sequencing. However, our model could be tested in larger future studies with long-read sequencing technology (Glinos et al. 2021).

Thus, we investigate common splice-regulatory variants (ψQTLs) as potential modifiers of penetrance of rare alleles in their target exons. Across different datasets, we have demonstrated and replicated the result that high-penetrance haplotype configurations of rare alleles and ψQTLs alleles are depleted. These findings emphasize the importance of alternative splicing as one of the many processes that regulate human traits, and suggest that splicing is involved in variable penetrance of coding variants.

Through this research, we derived a novel approach for calculating the cumulative distribution function of the Poisson-binomial distribution, as well as a metric for evaluating a dataset’s deviation from an expected distribution or difference between two data sets (the comparison test). This method is well suited for very large datasets, and has further applications in genetic and non-genetic analyses where data is expected to follow the Poisson-binomial.

While we were able to detect a genome-wide signal of nonrandom combinations of splice-associated and coding alleles, it must be noted that finding evidence of modified penetrance in population cohorts is difficult, and requires very large sample sizes. This is particularly true on an individual gene level: Even in a dataset as large as TOPMed, which contains tens of thousands of donors, few genes have reasonable statistical power to detect depletion of high-penetrance haplotype configurations individually. Furthermore, the biologically and medically important genes where variant penetrance is of most interest are also highly constrained and depleted of functional genetic variation overall, further limiting the data to test for haplotype combinations in the general population.

An alternative approach is to study regulatory variation underlying modified penetrance in disease cohorts with well annotated disease-causing variants, linking haplotype patterns with phenotype variation between and within families. The Simons Simplex Collection had some limitations in this respect: most ASD-contributing rare variants are not known and the trait is highly polygenic, making it difficult to compare penetrance of variants in the same gene between families. Furthermore, in simplex families many causal variants are *de-novo*, but their total number is small for statistical analysis. In the future, large ASD studies with multiplex families may better capture ASD instances with heritable variant etiology. Furthermore, experimental validation, for example with genome editing, may be a fruitful approach.

Overall these results suggest that depletion of high-penetrance ψQTL - coding variant haplotypes is robust across many data sources and gene sets. However, the data does not sufficiently support the hypothesis that modified penetrance by genetically controlled splicing is a significant driver for ASD risk, but that may provide some protection in families with a known incidence of autism.

In conclusion, this study provides evidence that splice-regulatory alleles play a role in controlling the impact of rare coding alleles with putatively deleterious effects. Understanding the importance of these mechanisms will be crucial for building a holistic model of genetic contribution to human phenotypic variation. We hope that in the future the prognosis of individuals carrying rare variants will be informed by genomic context that extends beyond coding regions.

## Methods

### Data Sources

In this project, we utilize bulk RNA sequencing and WGS from the Genotype-Tissue Expression (GTEx) Project Version 8 (Consortium 2020), WGS from 19 cohorts included in the Trans-Omics for Precision Medicine Project freeze 8 (https://topmed.nhlbi.nih.gov/topmed-whole-genome-sequencing-methods-freeze-8) (Supplemental Table 2) and WGS from simplex families in the Simons Simplex Collection (SSC).

### GTEx PSI quantification and filtering

Percent spliced in (PSI) was calculated from GTEx V8 RNA-seq data. We limited our analysis to 18 tissues, which were chosen for their coverage of tissue diversity GTEx and their coverage of the most coding genes possible (Table S1). Exon PSI for protein-coding genes was quantified using the Integrative Pipeline for Splicing Analysis (IPSA),(Pervouchine et al. 2013; 2020) which was run on Google Cloud through Terra (https://github.com/guigolab/ipsa-nf). The ‘-`unstranded’` flag was used during the sjcount process. Exons were defined by the modified version of Gencode annotation v26 used in GTEx V8, which collapses genes with multiple isoforms to a single isoform per gene (https://storage.googleapis.com/gtex_analysis_v8/reference/gencode.v26. GRCh38.genes.gtf).

For downstream analyses, PSI data for each tissue was prepared by 1) removing exons with data available in less than 50% of donors and 2) removing exons with fewer than 10 unique values across all available donors (Table S1). These data were normalized for QTL mapping by randomly breaking any ties between two individuals with the same PSI at an exon, then applying inverse-normal transformation across all individuals. Filtered and normalized PSI calls were saved in BED format with start/end position corresponding to each gene’s transcription start side (TSS), which serves as a reference for where to define windows for QTL mapping. The gene containing each exon was included in the BED files for use with QTLtools’ group permutation mode.

### PSI Z-Score Analysis in GTEx

We compiled a list of all exons with sufficiently variable splicing in at least one GTEx tissue, as defined in the previous step, and saved the genomic coordinates of these exons in BED format. Rare variants (gnomAD AF < .01) that fell on variably spliced exons were extracted from GTEx WGS VCFs, and were subsequently filtered to variants that appeared less than 6 and greater than 1 time. Rare variant CADD scores and annotations with respect to the relevant gene were extracted as well. Some rare variants were annotated as ‘intronic’ because CADD v1.5 uses a different annotation that in rare cases does not correspond to gencode v26. Rare variant calls from exons represented disproportionately, either due to length or to high number of variants at the exon, were removed. Threshold for removing an exon was defined as Q3 + 1.5 * IQR, where Q3 is the third quartile of the number of rare variants per exon, where IQR is the interquartile range of the number of rare variants per exon. For all remaining variants, we computed the PSI Z-score of the individual that carried the variant at that specific exon, across all tissues where the exon was expressed and sufficiently variable. The PSI-Z score for a particular individual *i* at an exon *j* in tissue *k* is calculated as (ψ_ijk_ - μ_jk_)/σ_jk_, where ψ_ijk_ is an individual’s PSI level at a particular exon and tissue, and μ_j_ and σ_j_ are the mean and standard deviation of PSI for an exon *j* across all individuals with data available for that exon in tissue *k*. Importantly, we do not normalize PSI for this analysis, to preserve signal from exons with high PSI Z-scores.

### Primary ψQTL mapping, collapsing, and secondary ψQTL mapping

For each of the 18 GTEx V8 tissue groups, QTL mapping was run on every exon that passed filtering, using all genetic variants with an allele frequency greater than 5% within 1Mb of the gene’s transcription start site. We used QTLtools (Delaneau et al. 2017) run in grouped permutation mode, with groups defined by gene. This strategy controls for correlation between exons that are part of the same gene. 15 PEER factors recalculated from normalized PSI, 5 genetic principal components (PCs), as well as sex, WGS PCR batch, and sequencing platform were also included as covariates in the QTL model, as recommended in the GTEx V8 STAR methods.(Consortium 2020)

For every exon, we selected the most significant variant, and for every gene the most significant exon. We then compiled the ψQTL results across tissues to achieve a set of cross-tissue top ψQTLs. When a gene was significant across multiple tissues, we used the tissue where the effect size (ΔPSI score) was the highest. This process ensured that a gene was only included once in our final set of ψQTLs, and was labeled by one variant that is associated to splicing (sVariant).

Since the splicing of multiple exons within a gene is often correlated, we implemented an approach to identify additional exons whose splicing the sVariant is associated with. Consideration of multiple exons per gene is desirable because it increases the amount of genetic space where rare variant haplotypes can be identified. For each gene with a significant ψQTL, we ran a nominal QTLtools pass of just the sVariant against PSI of all other exons in the gene. We then considered secondary exons with a Bonferroni-corrected p<0.05 if QTL effect direction was the same as the top exon.

This procedure produced the final set of common variant-exon pairs used in all downstream analyses (10,901 sExons, across 5,198 sGenes). Haplotype calls from phased, filtered WGS datasets (see next section) were compiled by extracting rare variants that fell within sExons, and recording if the variant appeared on the same haplotype as the high inclusion or low inclusion ψQTL allele. (Code available at https://github.com/jeinson/mp_manuscript)

### WGS filtering across datasets

#### Genotype Tissue Expression Project (GTEx)

Read-aware Phased WGS data was used from all 838 samples included in GTEx v8. (Consortium 2020), (Supplementary Information Section 2.4) For use in haplotype calling, the following filters were applied 1) Variants were extracted with an allele frequency less than 0.005 in gnomAD, and singleton variants without read-backing to support their phase call were removed. 2) Samples from donors that did not self-identify as European American were removed. Since the ψQTL data from GTEx is based on 85% European Americans, the sVariants selected from these data may not capture allele frequencies and haplotype structures in other ancestries, and differing numbers of rare variants across ancestries might bias the results. 3) Haplotype calls from genes represented disproportionately, either due to length or to high number of variants at the gene, were removed. Threshold for removing a gene was defined as Q3 + 1.5 * IQR, where Q3 is the third quartile of the number of haplotypes per gene, where IQR is the interquartile range of the number of haplotypes per gene.

#### Trans-Omics for Precision Medicine Initiative (TOPMed)

Population-phased WGS data from donors of European-American ancestry were used from TOPMed, since this matches the population source of the sQTL data from GTEx (see above). To define individuals of European ancestry, we used the approach outlined in (Morris et al. 2019). Briefly, TOPMed samples were projected onto the first 20 principal components estimated from the 1000 Genomes Phase 3 (1000G) project (Auton et al. 2015) using FastPCA v2.0 (Galinsky et al. 2016). Only bi-allelic variants shared between the two datasets, and that passed a strict set of criteria (MAF >1%, minor allele count >5, genotyping call rate >95%, Hardy-Weinberg p-value >1×10^−6^) were used to calculate the principal components. Expectation Maximization (EM) (Chen and Maitra 2015) clustering was used to compute the probabilities of cluster membership and eigenvectors 1, 2, 5, 6 and 8 were selected for efficiently separating the individuals of White European and American ancestry (subpopulation codes CEU, GBR, FIN, CEU, IBS and TSI) from other ancestry groups. Finally, eight predefined clusters were chosen for EM clustering based on sensitivity analyses. This resulted in 52,426 TOPMed individuals clustering together with the 1000G CEU, GBR, FIN, CEU, IBS and TSI subpopulation, and they were termed of White ancestry. We kept 19 cohorts (<u>Supplemental Table 2</u>), and 49,542 individuals, filtering out the remaining cohorts which collectively contained less than 5% of all haplotypes.

To define rare coding variants for downstream analysis, we extracted SNPs and small indels with more than 1 and 10 or fewer occurrences; singletons were removed due to unreliable population-based phasing. To account for unusually long genes, and genes with an unusually high number of rare variants, we applied the same filtering procedure as step 3 from the GTEx analysis to produce a final set of rare variant haplotypes.

#### Simons Simplex Collection (SSC)

Phased WGS data was used from 2,380 families. Simplex families consist of a proband child diagnosed with Autism Spectrum Disorder (ASD), an unaffected sibling, and two unaffected parents (Turner et al. 2016). We genotype the SSC whole-genome data set (An et al. 2018; Ruzzo et al. 2019; Yoon et al. 2021) using the transmission mode of our Multinomial Genotyper (Iossifov et al. 2012) that produces only high-quality mendelian family genotypes. The whole-genome sequence and the genotype calls are available to qualified researchers through the Simons Foundation. In addition, we transmission-phased the heterozygous variants on a per-variant basis when possible, using the genotypes of both parents. Since this method is accurate for singleton variants in probands, these were included in downstream analysis.

We additionally removed genes that contained an unusually high number of rare coding variants across parents, using the same outlier definition as in the previous two datasets. This set of variants post-filtering were considered in siblings and probands in downstream analyses.

#### Haplotype calling from phased genetic data and filtering

ψQTL-coding allele haplotypes were generated using a similar procedure across all three phased-resolved WGS datasets. First, all rare variants were extracted among sExons using the filters described above, considering variants that fell in primary and secondary sExons, taking account of the haplotype phase assignment. Then, the genotype of sVariants, and phase for heterozygous cases, was extracted from VCFs and haplotypes were labeled as high-penetrance (β = 1) and low penetrance (β = 0) according to our model for splice QTLs as a modifier of penetrance (Figure 1).

**Table 2:** Properties of 3 WGS datasets used in this study. Across all datasets, we extract rare variants that fall on primary and secondary sExons.

|  | **GTEx** | **TOPMed** | **SSC - Parents** |
| --- | --- | --- | --- |
| **N Donors** | 714 | 44,634 | 4,731 |
| **Phasing Method** | Population Based & Read backed phasing (SHAPEIT2(O'Connell et al. 2014) and PhASer (Castel et al. 2016)) | Population Phasing (Eagle) (Loh et al. 2016) | Phasing by transmission |
| **Singletons included** | Yes, in calls with RNA-seq read backing. Otherwise, no | No | Yes |
| **Rare variant allele frequency cutoff** | 0.5% MAF in gnomad. (No count cutoff due to the relative small size of the GTEx WGS dataset) | Appears 10 or fewer times (i.e. 0.0257% MAF) | Appears <= 3 times (i.e. 0.126% MAF) |

#### Test for depletion of regulatory haplotypes that increase penetrance

We sought to test the hypothesis that QTL-coding allele haplotype combinations are present in the population at frequencies that deviate from a baseline expectation, based on allele frequencies alone. Such a result could indicate high-penetrance haplotypes with deleterious variants being removed from the population by natural selection. The total number of high penetrance haplotypes arising from ψQTLs with varying allele frequencies can be modeled by the Poisson-Binomial distribution, which is a generalization of the binomial distribution. While a binomial describes the sum of *n* independent identically distributed bernoulli random variables, the Poisson-binomial describes the sum of *n* independent but non-identically distributed bernoulli random variables. Therefore, the distribution must be parameterized by a vector of probabilities of length *n*. While we could calculate P-values using a variety of methods that obtain the CDF of the Poisson-binomial, (Hong 2013) these methods all lack a way to quantify the magnitude of the effect size. Furthermore, they measure deviation from the null but do not allow comparison of two data sets (in our case, haplotypes carrying non-deleterious and deleterious coding alleles) Therefore, we developed the following procedure that approximates the Poisson-binomial CDF. This has the advantage of generating a quantifiable effect size for deviation from the null model, as well as corresponding confidence intervals.

Our procedure for approximating the Poisson-binomial, and subsequently testing for non-random occurrences of putative high-penetrance haplotypes, which we applied to each WGS dataset in this study, is as follows:

For each observation of a heterozygous coding allele that falls in a sExon, let *L* and *H* represent the low and high exon inclusion ψQTL haplotype respectively, and let *B* and *b* represent the coding variant reference and minor allele respectively. Here, we focus on rare variants, with our main interest being deleterious ones, and we here treat rare alleles as independent. Using variant phasing information, for a given haplotype *g*, we define an indicator function *β* which is set equal to 1, corresponding to putatively high-penetrance, if the coding allele falls on the highly included sExon, and 0 otherwise. The genotype of the major coding allele is irrelevant, and for rare variants *b/b* homozygotes are absent in practice.

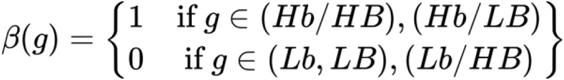

Next, we define an expectation function on *β*, under the null model where observing a high-penetrance and low-penetrance haplotype are equally likely. E[*β(g)*] is dependent on the heterozygosity of the ψQTL variant in an individual. Assuming independence of rare variants, if an individual is heterozygous for a ψQTL allele, the probability that an exonic variant will land in a high-penetrance configuration is 0.5. If an individual is homozygous for the ψQTL allele, the probability that the exonic variant will land in a high-penetrance configuration is dependent on the ψQTL’s allele frequency.

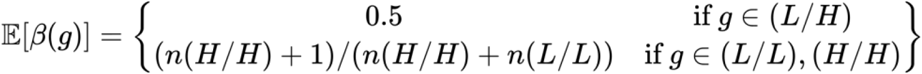

We define the expectation of observing a homozygous ψQTL allele as the proportion of high inclusion ψQTL homozygotes in the dataset, plus a pseudo-count, to avoid getting an expectation of 0 in datasets where the low inclusion allele is much more common. This method does not assume Hardy-Weinberg equilibrium for the ψQTL allele, but requires that the proportion of homozygotes for the two alleles be recalculated on each dataset. This approach was used for the GTEx and TOPMed analyses. Alternatively, the expectation of β under the null model can also be calculated as follows:

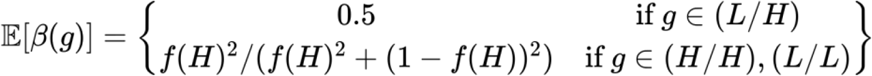

Where *f(H)* is the population frequency of the high exon inclusion ψQTL allele. We took this approach for haplotypes from SSC, where counting alleles across the whole dataset was infeasible due to the structure of the dataset, and used ψQTL allele frequencies from gnomad 3.0 (Karczewski et al. 2020).

The function *β* is evaluated across all individuals, sGenes, and rare variants in sExons in a dataset. The average observed deviation from the expected totals of high and low penetrance haplotypes (ε) is calculated as follows:

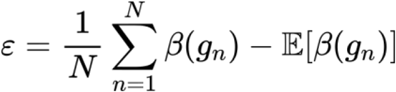

where *N* is the total number of considered haplotypes. ε can be interpreted as the effect size of depletion/enrichment of high-penetrance haplotypes in the dataset such that ε < 0 would indicate a depletion of high-penetrance haplotypes.

We quantify the significance of ε by bootstrapping all haplotypes, generating 95% confidence intervals and drawing two-sided empirical *P-*values as

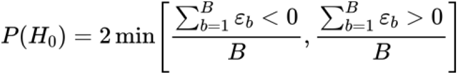

where *B* is the total number of bootstraps. In practice, we found that 1,000 bootstraps was enough to accurately approximate the Poisson-binomial distribution, while managing runtime.

Although the test was designed for counts of haplotypes, this approach is generalizable to any system that can be modeled by a Poisson-binomial distribution. Therefore, to benchmark our test, we simulated data from several theoretical allele frequency distributions by sampling from beta distributions with various shape parameters, including one distribution where its parameters were estimated direction from our set of ψQTLs from GTEx using the method of moments estimator (Figure 3, Supplemental Figure 4). We found that our bootstrapping procedure accurately approximated the Poisson-binomial distribution for all inputs tested. However, the magnitude of ε - but not direction - is dependent on the shape of the theoretical allele frequency distribution, so comparing magnitudes of ε across distinct datasets should be done with caution. The accuracy of our method increased with larger sample sizes. Therefore, we recommend using this approach when handling data where N > 1,000 (Supplemental Figure S4).

As an extension to this procedure, we can also conveniently calculate the significance of a difference in ε between two similar datasets *A* and *B*, for example, between haplotypes where the rare variant is putatively deleterious vs. haplotypes where the rare variant is non-deleterious:

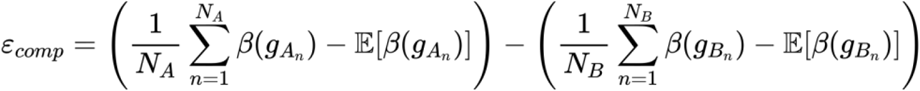

We then apply the bootstrapping procedure as in the standard case, and draw P-values accordingly. The corresponding P-value from this procedure is referred to as the “comparison test” in the main text.

This test is implemented in the STatististic for Modified PENetrance (STAMPEN) R package that is available to download here (https://github.com/jeinson/stampen)

## Data Availability

All code used to perform analyses and generate figures is available at https://github.com/jeinson/mp_manuscript. Qualified researchers requiring data access can apply for GTEx, and TOPMed data through dbGaP, and SSC data through the Simons foundation. We include a function to generate simulated data in the stampen R package (https://github.com/jeinson/stampen). PSI and ψQTLs from GTEx v8 can be download from the repository for (Einson et al. 2022) at https://zenodo.org/record/7275062#.Y9gc0OzMJf0

## Acknowledgements

J.E. thanks members of the Lappalainen lab for thoughtful discussions and feedback throughout this project.

Molecular data for the Trans-Omics in Precision Medicine (TOPMed) program was supported by the National Heart, Lung and Blood Institute (NHLBI). Whole genome sequencing (WGS) for the Trans-Omics in Precision Medicine (TOPMed) program was supported by the National Heart, Lung and Blood Institute (NHLBI). Core support including centralized genomic read mapping and genotype calling, along with variant quality metrics and filtering were provided by the TOPMed Informatics Research Center (3R01HL-117626-02S1; contract HHSN268201800002I). Core support including phenotype harmonization, data management, sample-identity QC, and general program coordination were provided by the TOPMed Data Coordinating Center (R01HL-120393; U01HL-120393; contract HHSN268201800001I) and TOPMed MESA Multi-Omics (HHSN2682015000031/HSN26800004).

Cohort specific acknowledgements for the 19 TOPMed cohorts used in this study are included in <u>Supplemental Table 2</u>. The content is solely the responsibility of the authors and does not necessarily represent the official views of the National Institutes of Health.

## Funding and Sequencing Center Information

1. Genome Sequencing for NHLBI TOPMed: Women’s Health Initiative (phs001237) was performed at Broad Institute Genomics Platform (HHSN268201500014C).
2. Genome Sequencing for NHLBI TOPMed: Genetic Epidemiology of COPD Study (phs000951) was performed at Northwest Genomics Center (3R01HL089856-08S1).
3. Genome Sequencing for NHLBI TOPMed: Atherosclerosis Risk in Communities Study VTE cohort (phs001211) was performed at Baylor College of Medicine Human Genome Sequencing Center (3U54HG003273-12S2 / HHSN268201500015C).
4. Genome Sequencing for NHLBI TOPMed: Framingham Heart Study (phs000974) was performed at Broad Institute Genomics Platform (HHSN268201600034I).
5. Genome Sequencing for NHLBI TOPMed: My Life, Our Future: Genotyping for Progress in Hemophilia (phs001515) was performed at Baylor College of Medicine Human Genome Sequencing Center (HHSN268201600033I).
6. Genome Sequencing for NHLBI TOPMed: Mount Sinai BioMe Biobank (phs001644) was performed at McDonnell Genome Institute (3UM1HG008853-01S2).
7. Genome Sequencing for NHLBI TOPMed: Cardiovascular Health Study (phs001368) was performed at Broad Institute Genomics Platform (HHSN268201600034I).
8. Genome Sequencing for NHLBI TOPMed: Multi-Ethnic Study of Atherosclerosis (phs001416) was performed at Broad Institute Genomics Platform (HHSN268201600034I, 3U54HG003067-13S1).
9. Genome Sequencing for NHLBI TOPMed: Coronary Artery Risk Development in Young Adults (phs001612) was performed at Baylor College of Medicine Human Genome Sequencing Center (HHSN268201600033I).
10. Genome Sequencing for NHLBI TOPMed: Mayo Clinic Venous Thromboembolism Study (phs001402) was performed at Baylor College of Medicine Human Genome Sequencing Center (3U54HG003273-12S2 / HHSN268201500015C).
11. Genome Sequencing for NHLBI TOPMed: Lung Tissue Research Consortium (phs001662) was performed at Broad Institute Genomics Platform (HHSN268201600034I).
12. Genome Sequencing for NHLBI TOPMed: The Vanderbilt University BioVU Atrial Fibrillation Genetics Study (phs001624) was performed at Baylor College of Medicine Human Genome Sequencing Center (3UM1HG008898-01S3).
13. Genome Sequencing for NHLBI TOPMed: Vanderbilt Genetic Basis of Atrial Fibrillation (phs001032) was performed at Broad Institute Genomics Platform (3R01HL092577-06S1).
14. Genome Sequencing for NHLBI TOPMed: Hispanic Community Health Study - Study of Latinos (phs001395) was performed at Baylor College of Medicine Human Genome Sequencing Center (HHSN268201600033I).
15. Genome Sequencing for NHLBI TOPMed: Severe Asthma Research Program (phs001446) was performed at New York Genome Center Genomics (HHSN268201500016C).
16. Genome Sequencing for NHLBI TOPMed: Massachusetts General Hospital Atrial Fibrillation Study (phs001062) was performed at Broad Institute Genomics Platform (3U54HG003067-12S2 / 3U54HG003067-13S1; 3U54HG003067-12S2 / 3U54HG003067-13S1; 3UM1HG008895-01S2).
17. Genome Sequencing for NHLBI TOPMed: Heart and Vascular Health Study (phs000993) was performed at Broad Institute Genomics Platform (3R01HL092577-06S1).
18. Genome Sequencing for NHLBI TOPMed: Groningen Genetics of Atrial Fibrillation Study (phs001725) was performed at Baylor College of Medicine Human Genome Sequencing Center (3UM1HG008898-01S3).
19. Genome Sequencing for NHLBI TOPMed: Genetics of Cardiometabolic Health in the Amish (phs000956) was performed at Broad Institute Genomics Platform (3R01HL121007-01S1).

J.E and TL were supported by NIH grants R01GM122924, R01MH106842. P.M. was supported by NIGMS grant R01GM140287. I.I. was supported by the Simons Center for Quantitative Biology at Cold Spring Harbor Laboratory, SFARI Grants SF497800, SF677963, SF666590, and the Centers for Common Disease Genomics grant (UM1 HG008901).Support for title page creation and format was provided by AuthorArranger, a tool developed at the National Cancer Institute.

## Supplemental Figures

**Figure S1.**
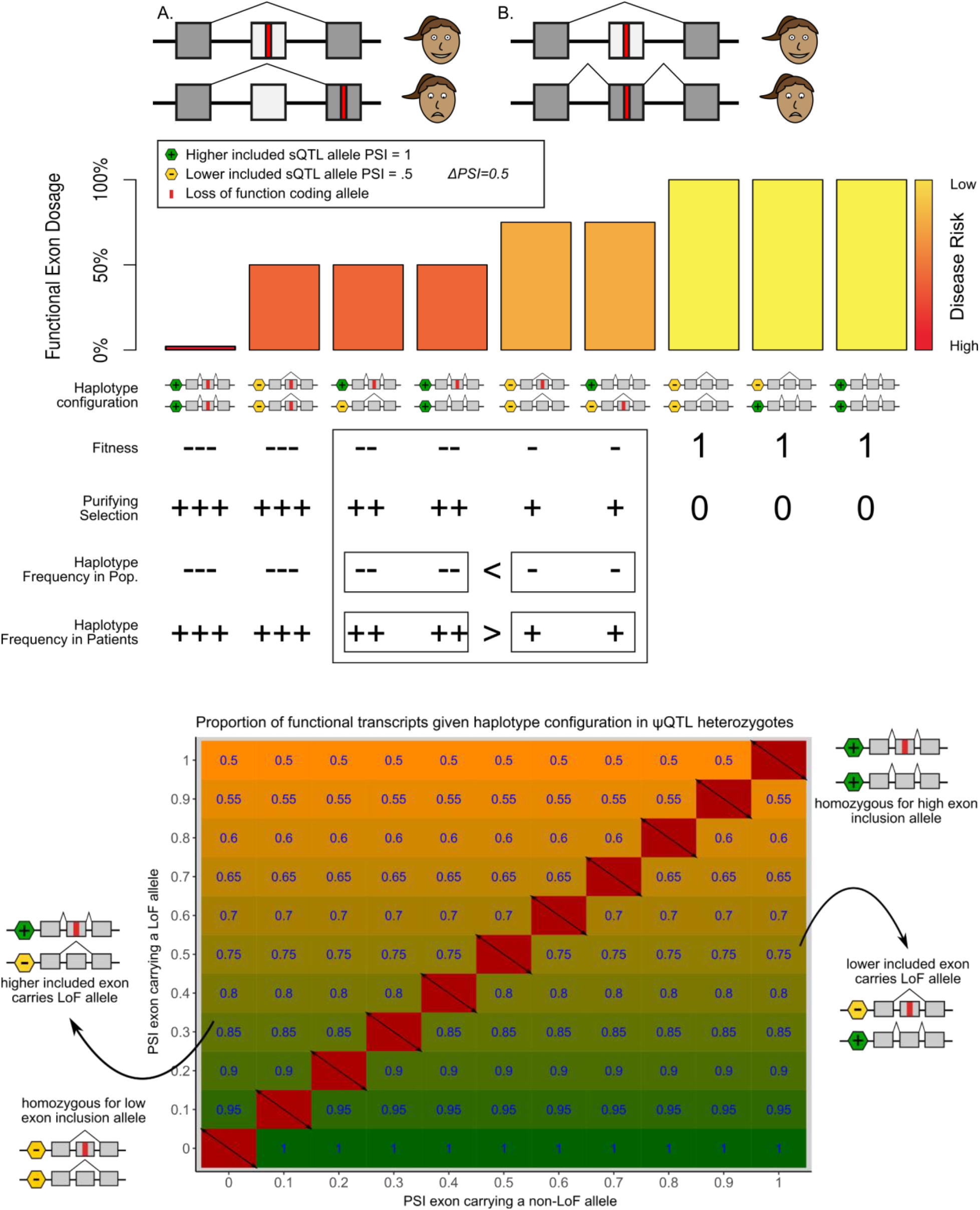
Splicing as a modifier of penetrance, in detail. Under the modified penetrance model that we consider here, regulatory variation that alters the dosage effect that a loss-of-function variant has on a gene is the primary driver of incomplete penetrance of the LoF variant. In this project, we focus specifically on exon splicing as a driver of this phenomenon. Generally, we consider regulatory alleles in this model to be selectively neutral, which is likely to be the case for most common regulatory variants. In the top example, we present a scenario where two splicing isoforms for a particular gene exist, and their ratio is controlled by a ψQTL where one allele causes a target exon to be included 100% of the time the gene is expressed, and the other allele causes the exon to be skipped 50% of the time it is expressed. If by chance, an individual carries a loss-of-function allele on the target exon (either by transmission or *de-novo* mutation), functional exon dosage is reduced to 75% if the loss-of-function variant lands on lower included haplotype. The functional dosage is further reduced to 50% if the loss-of-function variant lands on the higher included haplotype. In this example, it is important to note that loss of functional gene dosage is driven only by the haplotype carrying a loss-of-function allele, and its ψQTL allele being a potential modifier of this. The other haplotype is fully functional and its sQTL allele is irrelevant. This is a subtle but pertinent distinction between the eQTL as a modifier of penetrance hypothesis (Castel et al. 2018), where the LoF haplotype is considered non-functional, and the the non-LoF haplotype is responsible for maintaining normal downstream function as modified by its eQTL allele. All assumptions about haplotype frequency in the population and haplotype frequency in diseased patients are the same across the two models. In the lower figure, we generalize the model to include ψQTLs of all effect sizes. For heterozygotes, the upper left corner of the plot represents putative high-penetrance haplotypes, and the lower right corner represents putative low-penetrance haplotypes. For ψQTL homozygotes, deleterious or non-deleterious haplotype designations depend on the PSI of the alternative ψQTL allele. At the population level, we hypothesize that purifying selection acts more strongly against high-penetrance haplotype combinations. However, we do not account for quantitative changes in functional dosage as they are likely to be highly gene-specific and mostly unknown.

**Figure S2:**
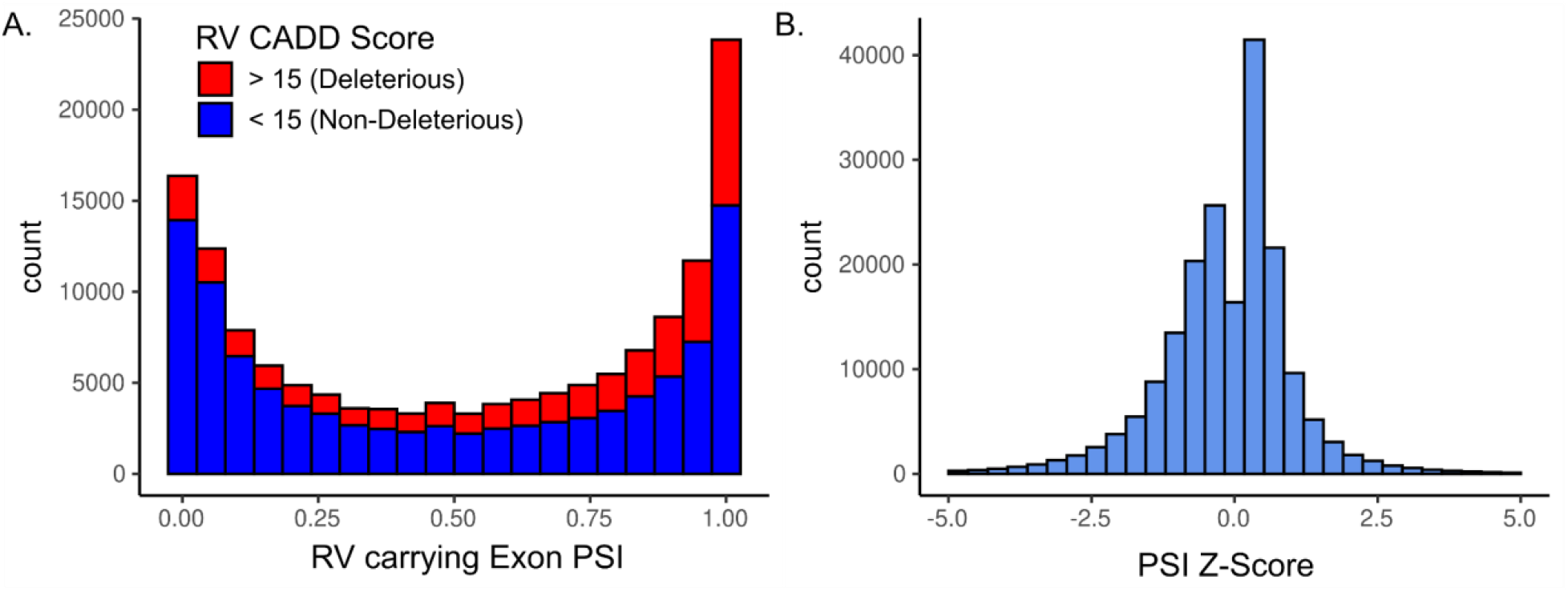
PSI among exons carrying a rare variant and PSI Z scores. **A**. Distribution of percent spliced in (PSI) scores of all exons with sufficient variability across individuals in GTEx that carry rare variants. Colors indicate CADD score of the rare variant. In general, variants on more highly included exons are assigned a higher CADD score. **B**. PSI Z-scores are generated by fitting a normal distribution to PSI levels across GTEx individuals for a particular exon. For each exon, the PSI Z-score is in reference to the splicing of the same exon in the same tissue across all other donors with RNA-seq data available for that tissue.

**Figure S3:**
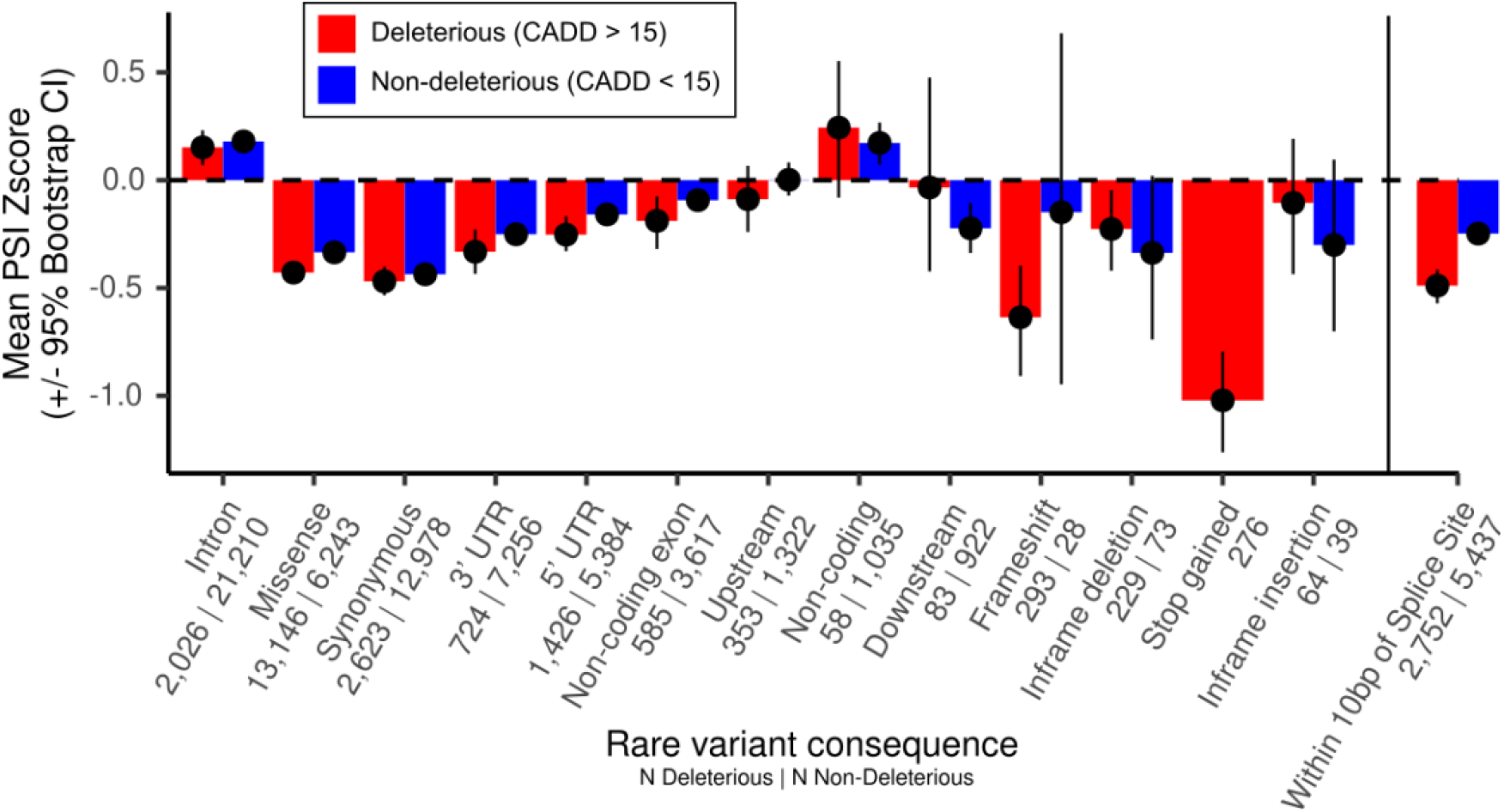
Mean Z-score (+/- 95% bootstrap CI) across annotations. The number of rare alleles with deleterious and non-deleterious CADD designations respectively are printed beneath each rare variant annotation. When data is available for an individual in multiple tissues, we calculated the mean Z-score. When collapsing across tissues and viewing by annotation, we see that deleterious alleles are depleted in most annotation classes as well. Some variants may be annotated as “intronic” even though their loci are labeled as exonic in the annotation used in the rest of the study.

**Figure S4:**
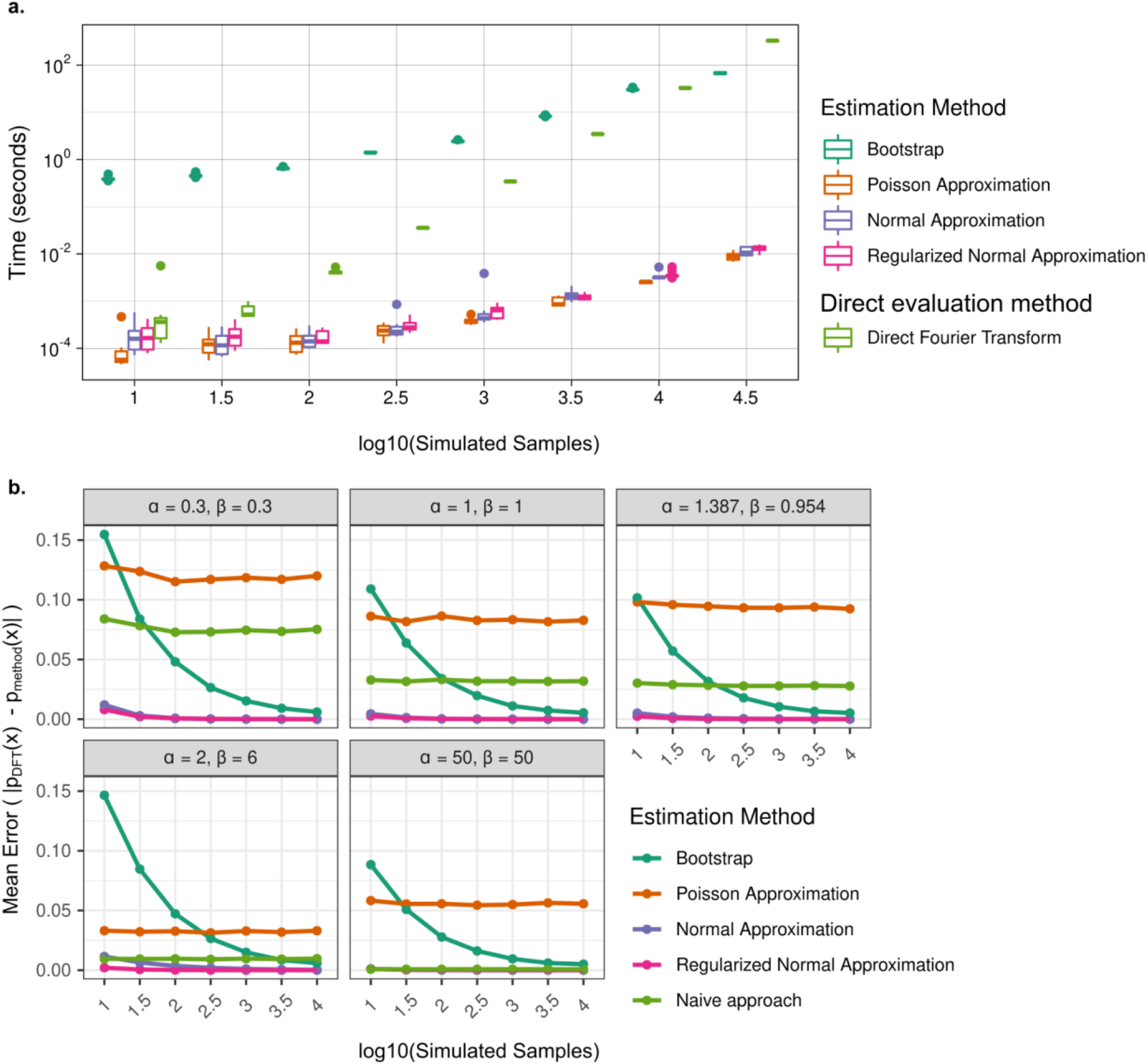
Runtime and Accuracy benchmark of the Bootstrapped Poisson-Binomial. For all benchmarking analyses, we compare our method, which approximates the cumulative distribution function (CDF) of the Poisson-binomial distribution using a bootstrapping procedure, to four other methods included in the ‘poibin’ R package. (Hong 2013) We use 5,000 bootstrap samples here, but we found that in general 1,000 bootstrap samples balanced accuracy and runtime. **a**. We measured the runtime to calculate the CDF of simulated datasets with uniform probability distributions. We found that the bootstrap method outperformed the Direct Fourier Transform (DFT) method for datasets with N > 10,000. DFT exceeded allocated memory for more than 10,000 samples, which we frequently encounter when analyzing real data. **b**. The bootstrap method performed more accurately with larger sample sizes, measured as the absolute difference between the estimation method and the DFT method. We tested across datasets with different distributions of *p*_*j*_, the vector of probabilities that define each binary observation. *p*_*j*_s were sampled from various beta distributions. The “naive approach,” for comparison, is a binomial test where *p* is the mean of *p*_*j*_.

**Figure S5:**
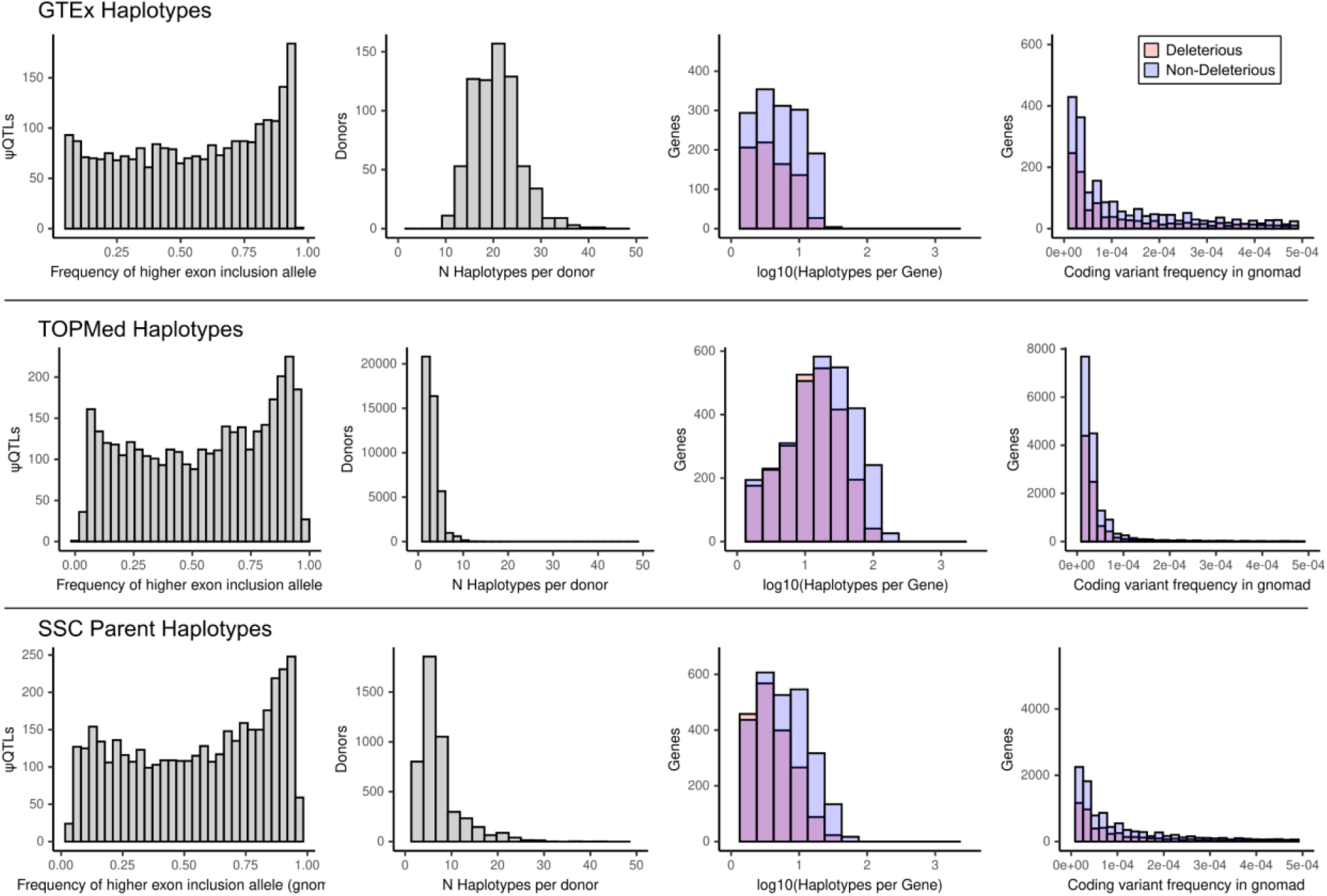
Summaries of haplotype calls across the 3 WGS datasets. In GTEx, TOPMed, and parents in the Simons Simplex Collection, we balanced sample size and allele frequency cutoffs to compile the best set of haplotype configurations. Across each dataset, we plot from left to right 1) the distribution of high exon inclusion ψQTL allele frequencies; 2) The number of haplotypes identified per donor, given the rare variant allele frequency cutoffs (see Table 2). The larger the dataset, the more stringent we can be for defining a ‘rare’ variant; 3) The number of haplotypes identified per gene; 4) The minor allele frequency in gnomad of all rare variants considered in the haplotype frequency analysis. Deleterious and non-deleterious refer to the CADD score designation (less than and greater than 15 respectively).

**Figure S6:**
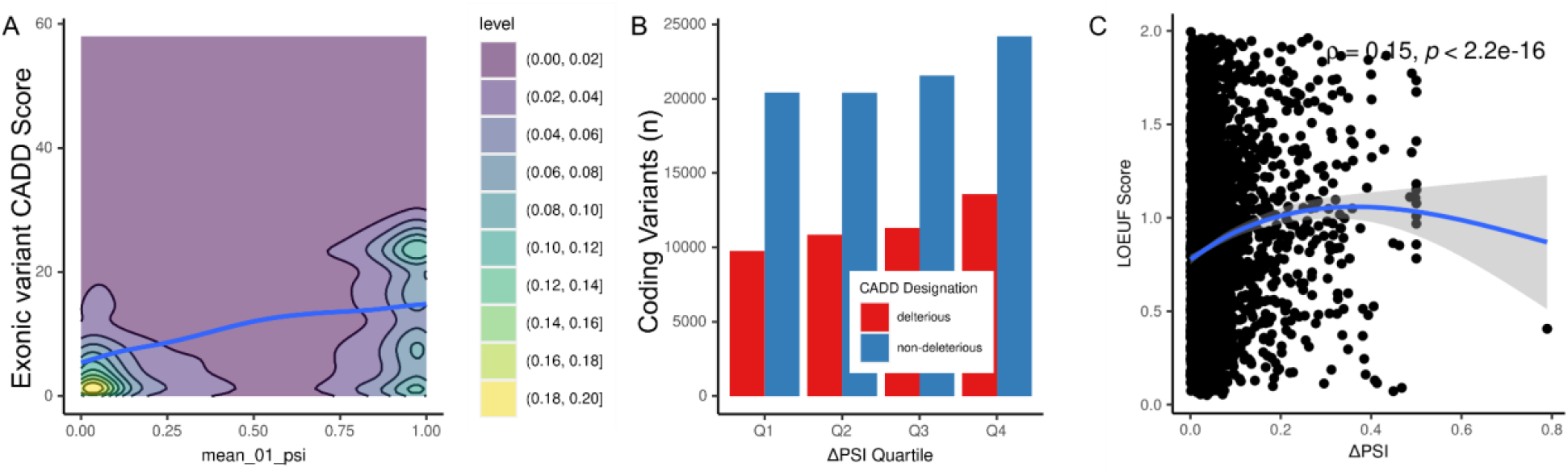
cSNP annotation counts in TOPMEd. **A**. More deleterious (higher CADD) variants tend to fall on exons with higher baseline PSI. **B**. Haplotypes grouped by ΔPSI Quantile. More rare variants, both deleterious and non-deleterious, appear at exons with larger effect size sQTLs. **C**. Genes that are tolerant to loss-of-function variants (high LOEUF) have ψQTLs with a higher effect size (ΔPSI).

**Figure S7:**
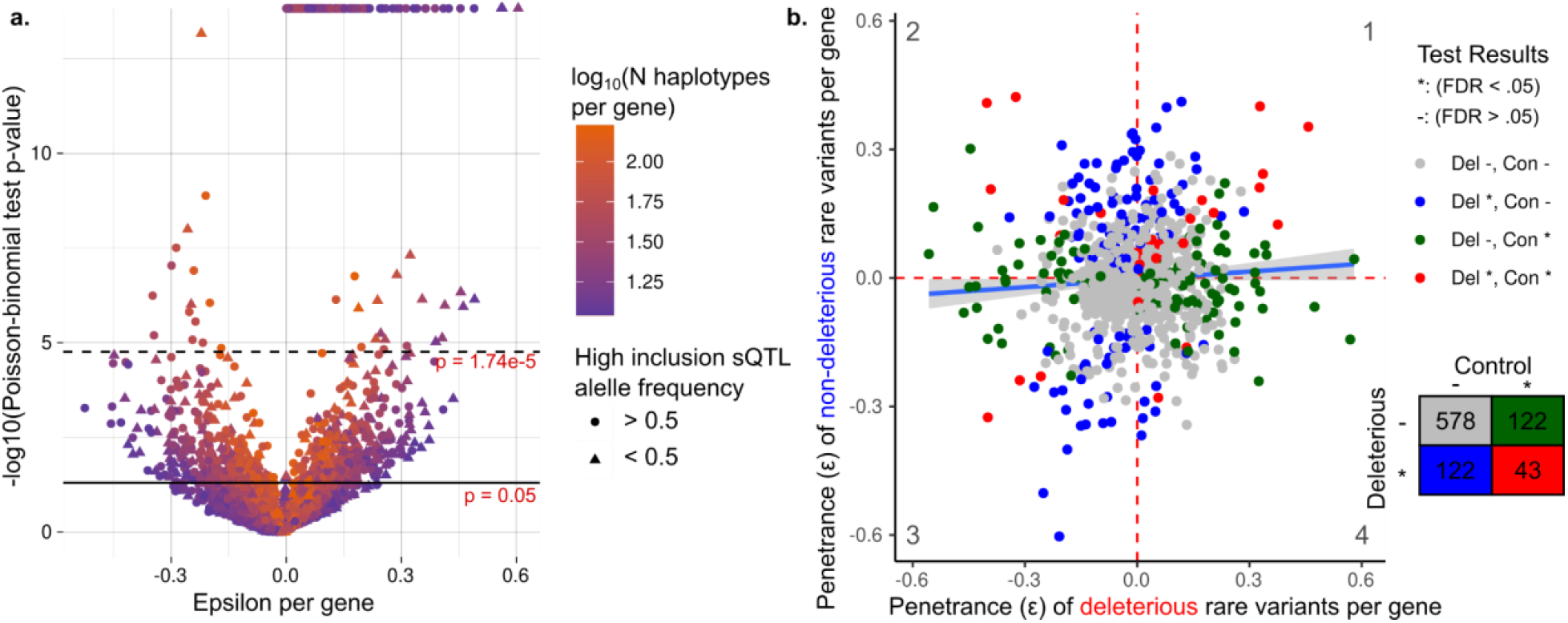
Gene by gene analysis in TOPMEd. **a**. Depletion of high-penetrance haplotype observations on a gene-by-gene basis in TOPMed. For each gene with more than 10 observed haplotypes across donors, we test if any genes or classes of genes are driving the overall pattern of high-penetrance haplotype depletion. Each point represents a single gene. **b**. Comparison of haplotype deviation between deleterious and non-deleterious rare coding variants, among genes with greater than 10 haplotypes in both categories. Under a model where highly penetrant deleterious cSNPs are depleted in the population, we expect more blue-labeled genes in the third quadrant.

**Figure S8:**
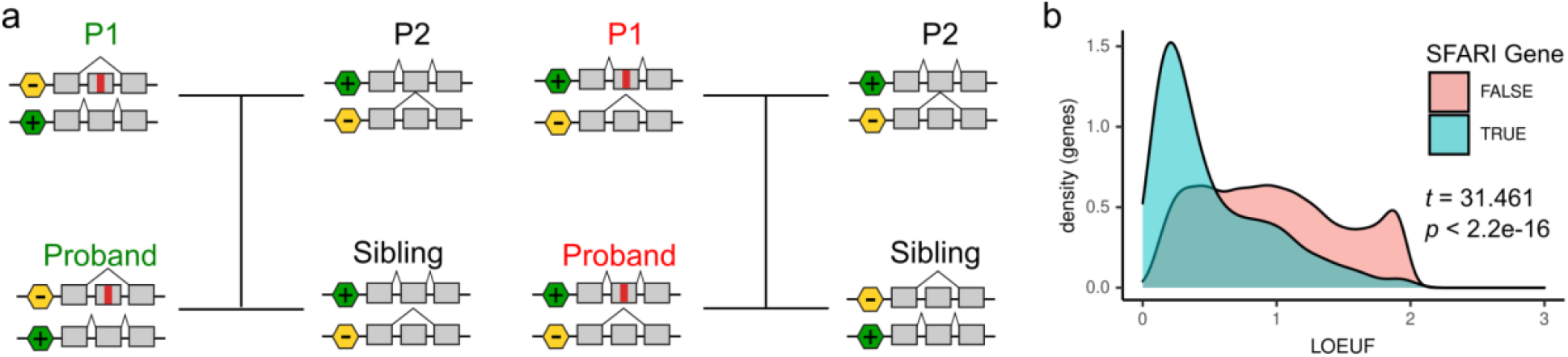
Transmission patterns of splicing haplotypes. **a**. When a parent carries a exonic variant in a putative low (green text) or high (red text) penetrance haplotype configuration, they will almost always transmit it to a child in the same haplotype configuration. **b**. Distribution of LOEUF scores among genes identified as relevant to Autism Spectrum Disorder, by SFARI Gene. ASD genes have significant depletion of predicted loss-of-function variants in general.

## Supplemental Tables

**Supplementary Table 1:** GTEx Tissues utilized for ψQTL calling, and the number of exons pre and post filtering.

| Tissue | N Exons per tissue pre-filtering | N Exons per tissue post-filtering | Percent usable | Genes covered per tissue |
| --- | --- | --- | --- | --- |
| Adipose_Subcutaneous | 260,800 | 29,180 | 11.19% | 8,585 |
| Artery_Tibial | 253,109 | 27,453 | 10.85% | 8,127 |
| Brain_Cerebellum | 239,928 | 36,095 | 15.04% | 8,605 |
| Brain_Cortex | 240,439 | 26,121 | 10.86% | 7,857 |
| Brain_Nucleus_accumbens_basal_ganglia | 247,074 | 26,372 | 10.67% | 7,998 |
| Cells_Cultured_fibroblasts | 230,752 | 28,486 | 12.34% | 8,479 |
| Cells_EBV.transformed_lymphocytes | 220,547 | 37,837 | 17.16% | 9,291 |
| Colon_Transverse | 231,647 | 29,066 | 12.55% | 8,630 |
| Esophagus_Mucosa | 245,627 | 26,721 | 10.88% | 7,984 |
| Liver | 224,469 | 21,605 | 9.62% | 6,283 |
| Lung | 265,555 | 34,585 | 13.02% | 9,387 |
| Muscle_Skeletal | 240,921 | 22,664 | 9.41% | 6,788 |
| Nerve_Tibial | 261,375 | 30,771 | 11.77% | 8,783 |
| Pituitary | 259,310 | 32,795 | 12.65% | 8,774 |
| Skin_Sun_Exposed_Lower_leg | 259,438 | 29,570 | 11.40% | 8,588 |
| Spleen | 241,122 | 30,379 | 12.60% | 8,277 |
| Thyroid | 266,364 | 30,035 | 11.28% | 8,586 |
| Whole_Blood | 236,866 | 23,135 | 9.77% | 6,039 |

**Supplemental Table 2: TOPMed cohorts utilized and number of samples from each cohort**

[Supplemental Table 2.xlsx](https://docs.google.com/spreadsheets/d/1QxM_m6kod8p1iWOjHOOt-tROOcLzJH0L/edit#gid=1060385474)

